# Structures and membrane interactions of native serotonin transporter in complexes with psychostimulants

**DOI:** 10.1101/2022.09.03.506477

**Authors:** Dongxue Yang, Zhiyu Zhao, Emad Tajkhorshid, Eric Gouaux

## Abstract

The serotonin transporter (SERT) is a member of the SLC6 neurotransmitter transporter family that mediates serotonin reuptake at presynaptic nerve terminals. SERT is the target of both therapeutic antidepressant drugs and illicit psychostimulant substances such as cocaine and methamphetamines, which are small molecules that perturb normal serotonergic transmission by interfering with serotonin transport. Despite decades of studies, important functional aspects of SERT such as the oligomerization state of native SERT and its interactions with potential proteins remain unresolved. Here we develop methods to isolate SERT from porcine brain (pSERT) using a mild, non-ionic detergent, utilize fluorescence-detection size-exclusion chromatography to investigate its oligomerization state and interactions with other proteins, and employ single-particle cryo-electron microscopy to elucidate the structures of pSERT in complexes with methamphetamine or cocaine, providing structural insights into psychostimulant recognition and accompanying pSERT conformations. Methamphetamine and cocaine both bind to SERT central site, stabilizing the transporter in an outward open conformation. We also identify densities attributable to multiple cholesterol or cholesteryl hemisuccinate (CHS) molecules, as well as to a detergent molecule bound to SERT allosteric site. Under our conditions of isolation, we find that pSERT is best described as a monomeric entity, isolated without interacting proteins, and is ensconced by multiple cholesterol or CHS molecules.

**Significance:** The serotonin transporter (SERT) is the target of antidepressants and illicit psychostimulants. Despite its importance in the nervous, cardiovascular and gastrointestinal systems, there is no direct knowledge of SERT’s oligomerization state(s) and interactions with other proteins. Here, we develop methods to isolate porcine SERT (pSERT) from native brain tissue in the presence of a mild, non-ionic detergent, and investigated its properties by biochemical, structural and computational methods. We show how cocaine and methamphetamine exert their pharmacological effect on SERT, binding to a site halfway across the membrane-spanning region of the transporter, stabilizing pSERT in an outward open conformation. pSERT is best described as a monomeric entity, requiring neither oligomerization or additional proteins for its structure or function.

## Introduction

Serotonin is a neurotransmitter that modulates multiple fundamental brain functions including memory, learning, sleep, pain, mood, and appetite (1). The serotonin transporter (SERT) removes serotonin from synaptic, perisynaptic and extracellular regions by harnessing the energy from sodium and chloride transmembrane gradients, diminishing local serotonin concentrations and thus terminating serotonergic neurotransmission (2, 3). Congruent with the crucial roles of serotonergic signaling in neurophysiology, dysfunction of SERT has profound consequences and is associated with neurological diseases and disorders, including Parkinson’s disease, seizures, depression, epilepsy, and attention deficit hyperactivity disorder (2, 4).

SERT is a member of the large neurotransmitter transporter family, also known as neurotransmitter sodium symporters (NSSs), a subfamily of the SLC6 transporters. In eukaryotes, additional members of the NSS family include transporters for norepinephrine (NET), dopamine (DAT), glycine (GlyT1 and GlyT2), and γ-aminobutyric acid (GAT), as well as for betaine and creatine (2), while in bacteria, homologs include LeuT (5) and MhsT (6). NSSs typically harbor 12 transmembrane helices organized topologically into two inverted repeats that, in turn comprise a conserved three-dimensional fold known as the LeuT fold (5). Substrate transport by NSSs can be described by an alternating access mechanism in which the substrate is translocated from extracellular to intracellular spaces (7, 8), a mechanism that has gained substantial support from a range of biochemical, biophysical and structural studies (4, 9–12).

SERT is a longstanding pharmacological target for antidepressant drugs (13), as well as the site of action for psychostimulants such as cocaine, amphetamine, and methamphetamine (14). The therapeutic utility of the drugs that act on SERT is a consequence of their specific action on SERT resulting in their relative lack of inhibition of the closely related DAT and NET transporters. In contrast, psychoactive drugs such as cocaine and amphetamines also inhibit or modulate the activity of DAT and NET, and as a consequence, they have pleotropic effects on the neurotransmitter reuptake systems, thus partially explaining their psychoactive and deleterious effects on neurophysiology and behavior (13). The potent and widely abused psychostimulants amphetamine and methamphetamine, or cocaine and its derivatives, act as substrates and promote transmitter efflux into the synaptic spaces or competitively inhibit the transport of neurotransmitters and lock the transporter in a transport-inactive conformation, respectively (2, 15). The x-ray structures of a transport-inactive *Drosophila melanogaster* DAT (dDAT) NSS transporter complexed with cocaine, amphetamine, or methamphetamine have revealed that illicit psychostimulants bind at the central substrate-binding site (16), yet studies with a functional transporter remained unresolved.

SERT interacts with a variety of intracellular scaffolding, cytoskeletal, anchoring, and signaling proteins. Examples include syntaxin1A, a vesicle fusion SNARE protein, which has been shown to interact directly with the amino terminus of SERT and regulate its surface expression level (17) and a neuronal nitric-oxide synthase (nNOS) that has a postsynaptic density of 95/discs-large/zona occludens (PDZ) domain interacts with the carboxy terminus of SERT, reducing its surface expression level and serotonin uptake capacity (18). While protein-protein interactions regulate SERT function and subcellular distribution, the extent to which they form stable complexes that can be biochemically isolated is not well understood. SERT also has cytoplasmic domains with numerous consensus sites for post-translational modification by protein kinases, phosphatases, and other interacting proteins that modulate its function and cellular distribution (19).

The oligomerization states of SERT and related NSSs have been a topic of debate for decades (20) and have been studied in the contexts of the plasma and organellar membranes (21, 22). Radiation inactivation and mutagenesis studies provided the first evidence for SERT oligomerization (23). Experiments with cross-linkers additionally suggested that rat SERT can form dimers and tetramers to varying degrees (24). Co-immunoprecipitation experiments (25–27), Förster resonance energy transfer (FRET) measurements (27–33), and fluorescence microscopy (32) (34) . Many of these studies were interpreted with an oligomerization model where the transporters form a variety of quaternary arrangements, ranging from monomers to multimers, differing to some extent depending on the specific NSSs (30, 32, 35–37). Membrane components such as phosphatidylinositol 4,5-bisphosphate (PIP2) and other lipids also have been implicated in the formation of NSS oligomers (38, 39), presumably via lipid interactions with the transmembrane helices, while psychostimulants such as methamphetamine and amphetamine have been shown to influence transporter oligomerization through an unknown mechanism (31, 40, 41).

Despite extensive experimental data from a broad range of biochemical, biophysical and computational studies (42, 43) that have been interpreted in terms of SERT oligomers, there has been no direct evidence for its oligomerization state based on the purified transporter isolated from a native source. Here, we develop methods to extract SERT from porcine brain tissue using the high-affinity 15B8 Fab and a mild non-ionic detergent, in the presence of methamphetamine or cocaine, allowing us to study the purified complex using fluorescence-detection size-exclusion chromatography (FSEC). We then carry out high-resolution, single-particle cryo-electron microscopy (cryo-EM) reconstructions, together with computational studies, to probe the conformation of psychostimulant-bound transporter and its interaction with lipids of the native cell membrane.

## Results

### Purification and cryo-EM structural determination of SERT

To isolate SERT, we exploited the 15B8 Fab (44), an antibody fragment that binds to a tertiary epitope of human SERT (hSERT), yet does not hinder the binding of ligands or the transport activity. We hypothesized that because porcine SERT (pSERT) and hSERT are closely related (*SI Appendix*, Fig. S1), the 15B8 Fab would also bind to pSERT and could serve as a powerful tool for immunoaffinity isolation of the transporter. We thus engineered the 15B8 Fab with a carboxy terminal mCherry fluorophore and an affinity tag (Fig. 1a).

**Fig. 1.**
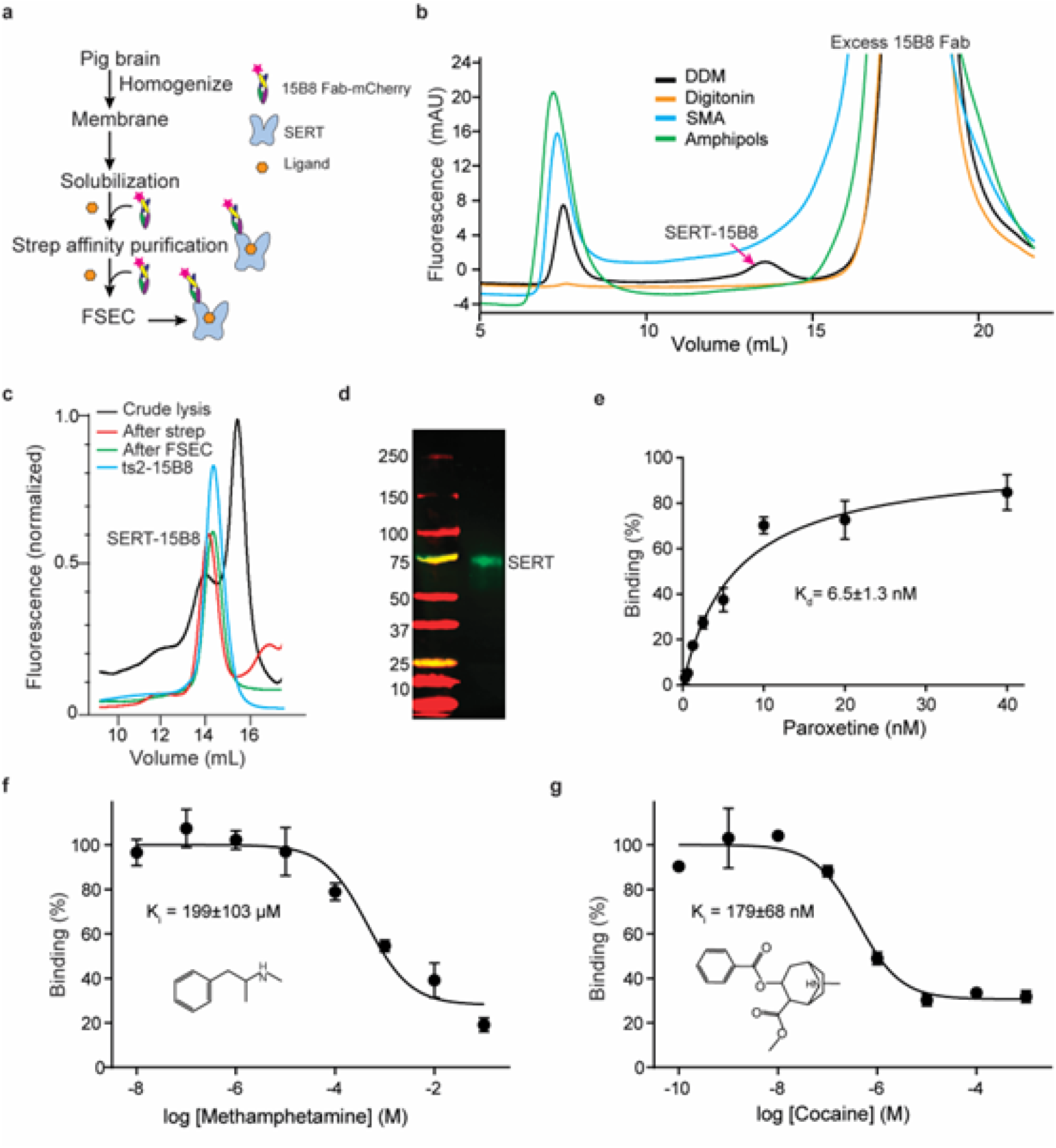
Purification and biochemical analysis of pSERT from native membranes. **a** Flow chart for pSERT purification. **b** FSEC profiles for screening of solubilization conditions. **c** Representative FSEC profile for pSERT in complex with the 15B8 Fab. **d** Western blot analysis of isolated pSERT after FSEC. The experiments were repeated two times with similar results. **e** Saturation binding of [^3^H] paroxetine to pSERT. **f** Competition binding of (+)-methamphetamine with [^3^H]paroxetine for pSERT. Symbols show the mean values derived from n=3 technical replicates. Error bars show the SEM. **g** Plots of competition binding of cocaine against [^3^H]paroxetine for pSERT. Data are means ± SEM.

To isolate pSERT from brain membranes, we explored different membrane protein solubilization conditions, aiming to extract the transporter under the mildest of conditions while retaining as much surrounding native lipids as possible. Solubilization in the presence of styrene-maleic acid (SMA) co-polymer (45) or the recently developed amphipols (46) (47), did not yield a measurable amount of native pSERT (Fig. 1b). We then examined the mild non-ionic detergents, digitonin or n-dodecyl-β-D-maltoside (DDM) together with cholesterol hemisuccinate (CHS), based on their utility in extraction of native (48) or recombinant hSERT (49), respectively. Surprisingly, a peak in the FSEC trace for the SERT-15B8 Fab- mCherry complex was only observed with the DDM/CHS mixture (Fig. 1b). We therefore utilized DDM/CHS in all subsequent studies. To isolate pSERT from brain tissue, we incubated the solubilized membranes with an excess of the 15B8 Fab-mCherry protein, as well as with saturating concentrations of either methamphetamine or cocaine. The transporter was purified by affinity chromatography, followed by FSEC, collecting fractions manually (Fig. 1c). Analysis of the isolated material by western blot revealed a band migrating with an apparent mass of 75 kDa (Fig. 1d), consistent with the estimated mass of pSERT. The well-resolved and symmetrical FSEC peak indicated that the purified pSERT 15B8 Fab-mCherry complex was monodisperse and best described as a single species. The elution volume with FSEC of pSERT 15B8 Fab complex was consistent with the recombinant ts2-15B8 complex (Fig. 1c), indicating that pSERT purified in the presence of DDM/CHS, is best described as a monomer.

To explore the function of pSERT, we then carried out saturation binding experiments using the high-affinity SSRI [^3^H]paroxetine for which we determined a dissociation constant (*K*_d_) of 6.5 ± 1.3 nM (Fig. 1e), in agreement with previous measurements (50). To characterize methamphetamine and cocaine binding with pSERT, competition experiments were performed, similarly employing [^3^H]paroxetine, and measuring inhibitory constants (*K*_i_) of 199 ± 103.4 µM (Fig. 1f) and 179 ± 68 nM (Fig. 1g), respectively, thus indicating that both methamphetamine and cocaine compete for [^3^H]paroxetine binding, consistent with binding of the psychostimulants to the central site. The potencies of methamphetamine and cocaine on pSERT differ by a factor of ∼1,000, in accord with previous studies, whereas on DAT the difference is only a factor of ∼10 (51), underscoring the differences in residue composition and plasticity of the central binding pockets of SERT and DAT.

From one porcine brain we obtained ∼20 μg of purified protein in a volume of 200 μL, which was sufficient for visualizing particles on continuous carbon-coated grids under cryogenic conditions. We then collected single-particle cryo-EM data sets and carried out reconstructions of the methamphetamine- or cocaine-bound SERT complexes, obtaining density maps that extended to approximately 2.9- and 3.3-Å resolution, respectively (*SI Appendix*, Fig. S2-6, Fig. S7a-b and Table 1). Particle picking was employed with an elliptical ‘blob’ of sufficient size to capture pSERT oligomers. Thorough 2D and 3D classifications yielded only a single class for each data set in which pSERT was found as a monomeric entity, with no evidence of dimers or higher-ordered oligomers (*SI Appendix*, Fig. S2-3), consistent with the molecular size of SERT estimated by FSEC. Overall, the density maps are of sufficient quality to assign most of the amino acid side chains, to identify additional non-protein density within the central binding site as bound ligands, and to suggest the presence of cholesterol or CHS molecules surrounding the transporter transmembrane domains (Fig. 2).

**Fig. 2.**
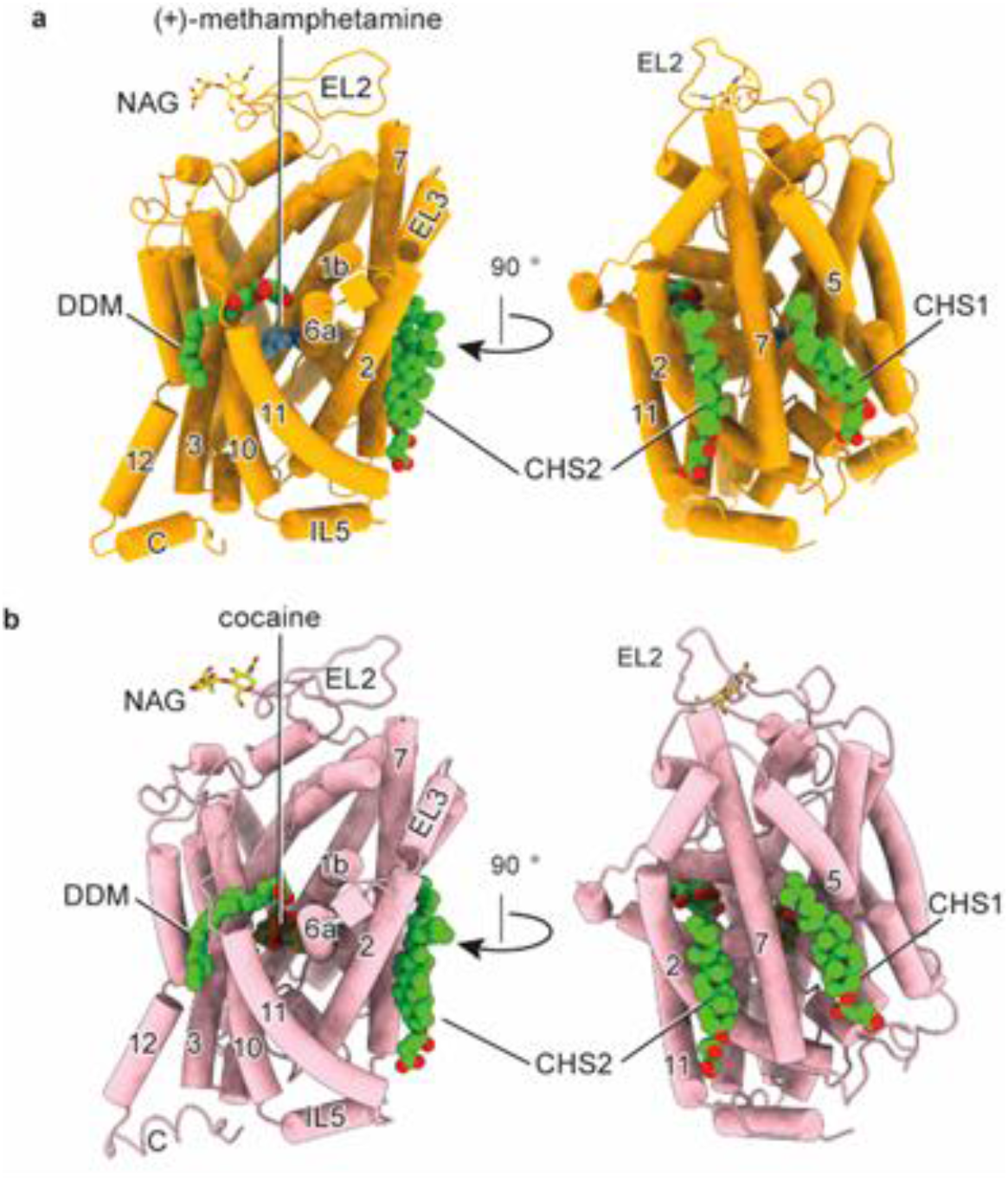
The cryo-EM structure of pSERT in complex with (+)-methamphetamine or cocaine, respectively. **a** Overall structure of the (+)-methamphetamine complex in the outward-open conformation, shown in cartoon representation. **b** Cartoon representation of the cocaine complex in the outward-occluded conformation. (+)-methamphetamine, cocaine, cholesteryl hemisuccinate (CHS), and n-dodecyl-β-D-maltoside (DDM) are shown in space- filling representations.

### Psychostimulant occupancy of the central site

Methamphetamine binds to the central site of the pSERT complex, adopting a similar binding pose to that observed in DAT (16), lodged between the aromatic groups of Tyr213 and Tyr132. The amine groups of methamphetamine interact with Ser475 and form hydrogen bonds with the carboxylate of Asp135 and the main chain carbonyl of Phe372, as seen with the hydrogen bonds formed between the amine group of methamphetamine and the equivalent Asp46 and Phe319 in DAT (16) (PDB code: 4XP6). The side chain of Phe378 forms edge-to-face aromatic interactions with the phenyl group of methamphetamine (Fig. 3a, *SI Appendix*, Fig. S7c). Further studies, at higher resolution, and complexes of methamphetamine with DAT, will be required to better explain the weaker binding of methamphetamine to SERT in comparison to DAT. Comparison of the positions of TM1, TM6, and the extracellular gate to the equivalent elements of serotonin bound outward-open complex (52) (Protein Data Bank (PDB) code: 7LIA, RMSD: 0.606) indicates that the SERT-15B8 Fab-methamphetamine complex adopts an outward-open conformation (Fig. 3b, *SI Appendix*, Fig. S7d), which is consistent with previous structural studies of dDAT in complex with methamphetamine (16).

**Fig. 3.**
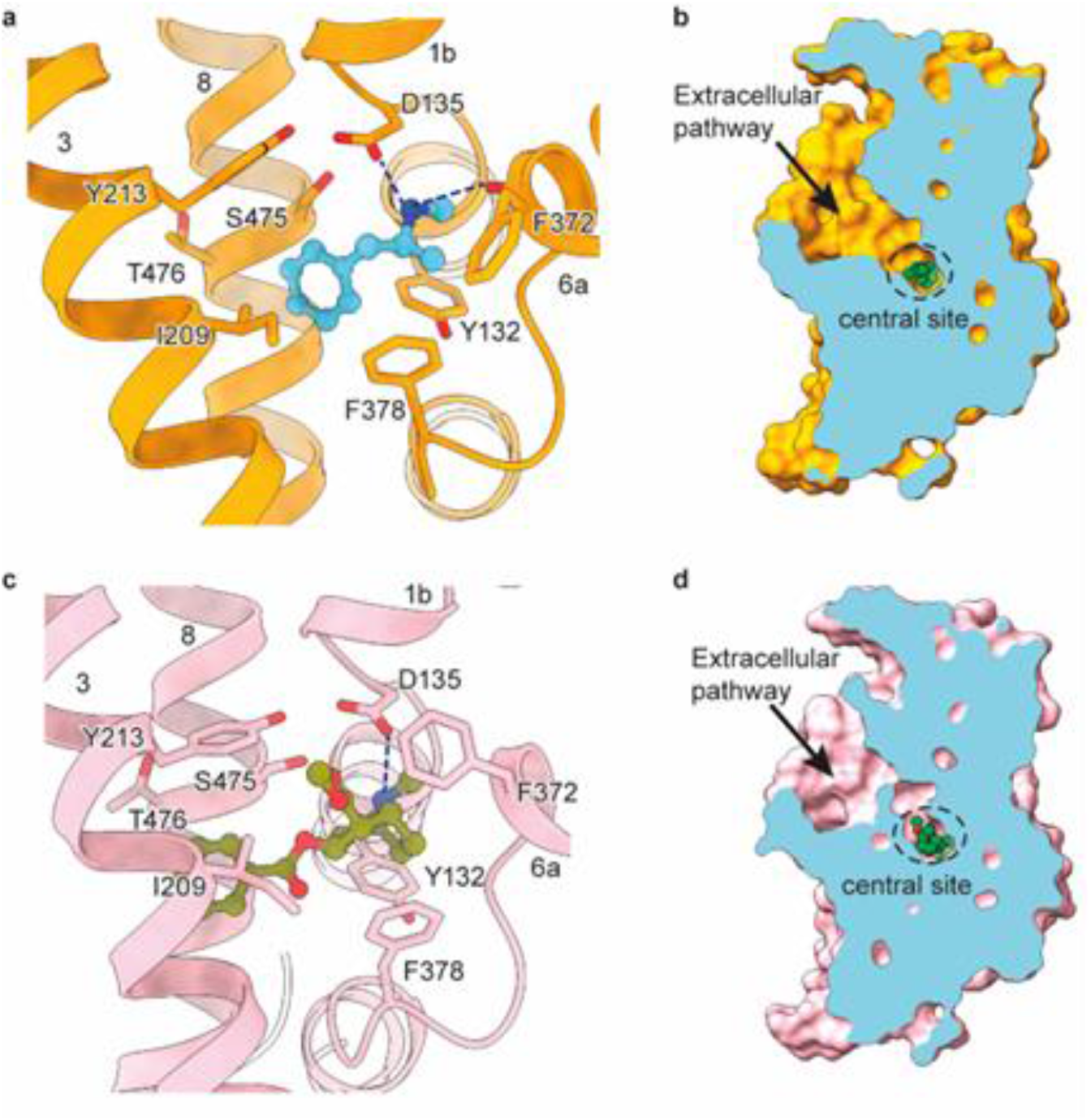
Ligands occupy the central site. **a** Close-up view of (+)-methamphetamine in the binding pocket with hydrogen bonds shown as dashed lines. **b** Slice view of pSERT in complex with (+)-methamphetamine. **c** Cocaine interactions within the central binding site. A hydrogen bond between cocaine and D135 is indicated with a dashed line. **d** Slab views of the extracellular cavity of pSERT in complex with cocaine.

The structure of the pSERT-cocaine complex displays an outward-open conformation, with cocaine occupying the entire central binding pocket with an overall similar pose to the dDAT-cocaine complex (16). The nearly complete filling of the volume of the central site by cocaine perhaps accounts for the increased affinity of SERT for cocaine compared to methamphetamine, reminiscent of how increasing the volume of serotonin via methylation can enhance ligand binding (52). The benzoyl moiety of cocaine is accommodated between TM3 and TM8, where it forms van der Waals interactions with Ile209, Tyr213, Phe378, and Thr476. The methyl ester group protrudes into the base of the extracellular vestibule and the tropane rings are bordered by Tyr132, Ala133, Asp135, Phe372, and Ser475. Interestingly, the side chain of Phe372 undergoes substantial displacement and moves further into the central site than seen in the dDAT complex (16). This reorganization translates into the formation of a ‘thin’ extracellular gate (53), blocking the release of cocaine from the central site and ultimately occluding the binding pocket, a conformation that was not visualized in the cocaine-bound dDAT structure (Fig. 3c, *SI Appendix*, Fig. S7e). Nevertheless, the overall structure of the SERT-15B8 Fab-cocaine complex is similar to 5-HT bound recombinant hSERT in its outward open conformation (PDB code: 7LIA, RMSD: 0.6 Å), with residue pairs that define the extracellular gate, Arg141-Glu530 and Tyr213-Phe372, 11.1 Å and 15.0 Å apart, measured from Cα to Cα, respectively (Fig. 3d, *SI Appendix*, Fig. S7f).

### Lipid binding sites surround the TMD

SERT is an integral membrane protein embedded in a complex membrane composed of phospholipids, sphingolipids, and cholesterol (54). SERT (54–56), NET (57, 58), DAT (59–61), and GlyT (62, 63), as well as some excitatory amino acid transporters (64) associate with cholesterol in brain tissues or in transfected cell lines. Cholesterol is implicated in a variety of biological processes, including membrane protein organization and compartmentalization within the membrane. It is also known to play a key indirect role in modulating neurotransmission via its effects on the activities of DAT (65) and SERT (54). Indeed, depletion of cholesterol from membranes affects the function of neurotransmitter transporters (65, 66). Previous molecular dynamics (MD) studies revealed six potential cholesterol binding sites of SERT, defined as CHOL1-6 (67). Bound cholesterol has been observed at the CHOL1 binding site in dDAT structures (16). The cholesterol analog, CHS, has been shown to bind at the CHOL2 binding site in dDAT (16, 68), as well as in hSERT (44), as well as at the CHOL3 binding site in hSERT (50). In investigating the interactions of cholesterol with pSERT, we carefully examined the density maps of methamphetamine- and cocaine-bound SERT complexes, and the quality of the density maps enabled the identification of CHS, or perhaps cholesterol, at the CHOL1 and CHOL2 binding sites in both structures (Fig. 4a-b, d-e), consistent with previous observations.

**Fig. 4.**
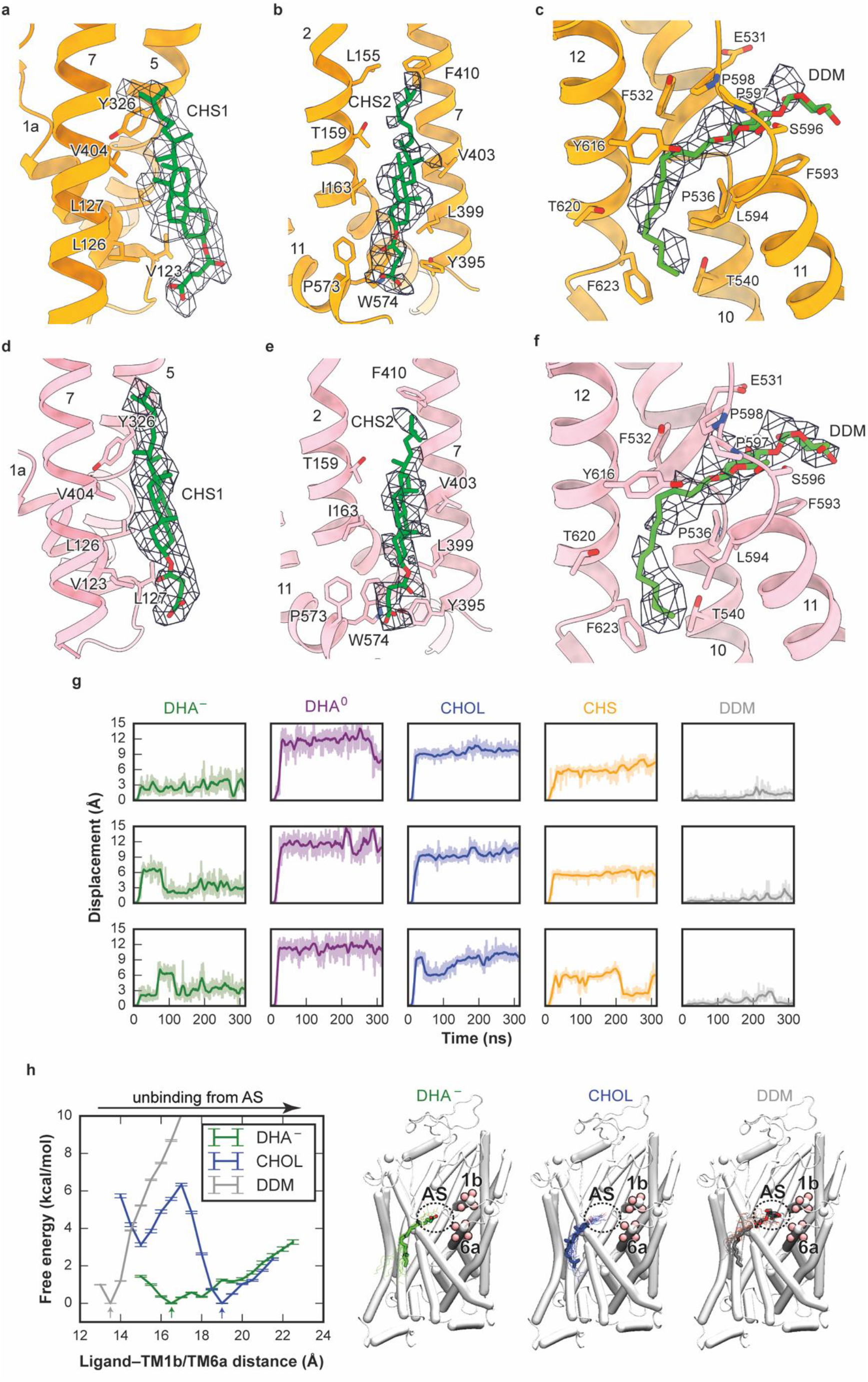
Cholesteryl hemisuccinate (CHS) and DDM binding in SERT. **a** and **d** Close-up views of CHS modeled at the junction of TM1, TM5, and TM7 interacting with multiple hydrophobic residues. **b** and **e**, CHS modeled at the junction of TM2, TM7, and TM11. **c** and **f**, DDM modeled at the allosteric site. **g** Time series of center-of-mass displacements of the ligands at the allosteric site (AS). Data for DHA^−^, DHA^0^, CHOL, CHS, and DDM are plotted in green, purple, blue, orange, and gray, respectively, and are shown for the three independent simulations in each case. Plots are smoothed using a sliding window of 1 ns. **h** Free energy profiles of DHA^−^, CHOL, and DDM binding to the allosteric site measured along the ligand- TM1b/TM6a distance, with molecular images showing each molecule in its most energetically favorable position (arrowed in left image). The ligand-TM1b/TM6a distance is measured as the center-of-mass distance between heavy atoms in the ligand and Cα atoms from TM1b and TM6a (residues 145-148 and 361-364, shown as pink spheres). The licorice representations represent DHA^−^, CHOL, and DDM in their most energetically favorable positions among all the examined ones (shown in line representations).

We identified a non-protein density in the allosteric site for both complexes (Fig. 4c, 4f), a binding site for a broad spectrum of ligands (50, 52, 69) (*SI Appendix*, Fig. S7g-i). Previously, the maltose headgroup of a DDM molecule has been modeled into this site, based on a cryo-EM map from heterologously-expressed hSERT (70). Because we isolated SERT from native tissue and the overall shape of the density also resembles the lipid molecule, we considered the possibility that the density feature might also be attributed to a lipid molecule. Because the local resolutions of the density maps within the allosteric site are not sufficient for unambiguous molecular identification, we used MD simulations to examine the binding of the most abundant lipid molecules, steroids, or fatty acids to the allosteric site. We performed all-atom MD simulations of the cocaine-pSERT complex with the allosteric site occupied by either docosahexaenoic acid (DHA), in charged or neutral forms (DHA^−^ or DHA^0^, respectively), cholesterol, CHS, or DDM, in triplicate simulation replicas for each molecular species (Fig. 4g, *SI Appendix*, Fig. S8a).

In all three independent simulations, DDM remained bound stably at the site (center-of-mass displacement ≤ 3 Å). DHA^−^ remained bound in one simulation, but in the other two, transiently moved away and then returned to its initial location. By contrast, in all simulations with neutral DHA^0^, cholesterol, or CHS, the ligand unbinds from its binding site within nanoseconds, as highlighted by large center-of-mass displacement (>5 Å). We further performed bias-exchange umbrella sampling simulations to calculate binding free energy profiles for DHA^−^, cholesterol, and DDM, verifying preferential binding of DDM, and DHA^−^ to a lesser extent, to the allosteric site, whereas cholesterol binding to this site appears to be accompanied with a large increase in free energy (Fig. 4h, *SI Appendix*, Fig. S8b-c). These results suggest that the allosteric site is likely occupied by DDM under our experimental conditions, yet it can also accommodate fatty acid binding. However, cholesterol binding to this site is energetically unfavorable. The biological role of lipid binding within the allosteric site awaits further elucidation.

## Discussion

Despite decades of experimental studies suggesting that monoamine transporters form oligomeric quaternary complexes (20), there has been no direct determination of oligomerization state for these proteins using purified native transporters isolated under mild conditions. In this study, we used immunoaffinity purification to isolate pSERT directly from brain membranes, proceeding to solve its cryo-EM structure in complex with the 15B8 Fab, and together with FSEC data, we show that pSERT, isolated in the presence of DDM/CHS, is best described as a monomer rather than as a dimer or a multimer (Fig. 1-2). Previous biochemical studies suggested that TM11 and TM12 form oligomeric interfaces in hSERT, and also suggested potential contributions by TM5 and TM6 (29). By contrast, the x-ray structure of SERT indicates that the kinked TM12 and the additional C-terminal helix protruding into the membrane preclude dimerization of SERT via a LeuT-like, TM12 interface (5). Furthermore, subsequent structural studies revealed that TM5 and TM6 are directly involved in the transporter’s conformational transitions between different functional states, and thus we speculate that their required flexibility is likely incompatible with the formation of dimers or multimers. Nevertheless, our study does not exclude the possibility of dimeric or multimeric arrangements of SERT that may be present in native membranes. Furthermore, because we carried out the isolation of pSERT in the presence of DDM/CHS, perhaps the loss of lipids, such as PIP2 (39), during membrane extraction and purification, yields exclusively monomeric transporter. Thus, further studies are needed to explore different methods of pSERT isolation and to investigate the roles that membrane constituents, such as PIP2, may play in oligomer formation and stability.

Prior structural studies of transport-inactive dDAT complexed with methamphetamine, cocaine, or their analogs provided a structural framework for showing how addictive psychostimulants stabilize the transporter in an outward open conformation (16). Here, we employed pSERT and single particle cryo-EM to study methamphetamine and cocaine binding. We find that compared to the corresponding dDAT structures, methamphetamine and cocaine have similar binding site locations and interactions at the central binding site of SERT. There are differences in the transporter gating residues, however, with cocaine occupying an outward open conformation of SERT, and with Phe372 rotating ‘inward’ to cover the tropane ring of cocaine, thereby blocking the release of the ligand from the central site, a conformational change not seen in the transport-inactive, cocaine-bound dDAT structures (Fig. 3).

Cholesterol is implicated in the function of SERT via direct cholesterol-protein interactions and modulates its function by enhancing substrate transport and antagonist binding (54). MD simulations show that cholesterol molecules are embedded in multiple sites of SERT (67), three of which have been confirmed by structural studies (16, 44, 50, 68). Here we discovered two cholesterol binding sites in pSERT. In addition, we also observed a non-protein density in the serotonin allosteric site (Fig. 4). We find that DDM is well accommodated into the experimental density and is stably bound as determined by MD simulations and free energy calculations. Meanwhile, this region of TM10, TM11, and TM12 has been indicated to be a potential lipid-binding site, and our MD simulations suggest that DHA also can be accommodated within this site. Conclusive determination of the native lipid molecules that bind to the allosteric site and their potential functional roles await further studies.

Using biochemical analysis and cryo-EM, we observed that in the presence of mild non-ionic detergents, the native, mammalian pSERT is isolated as a monomer, without interacting proteins, but bound with multiple cholesterol and lipid molecules. We investigated methamphetamine and cocaine binding to pSERT and show that both ligands occupy the central site, where they are involved in numerous interactions with surrounding residues. Moreover, despite extensive 3D classification in the single particle analysis, we find that pSERT occupies a single conformation when bound with either ligand, in contrast to the ensembles of conformations found in the presence of either ibogaine (44) or serotonin (52). Our studies of pSERT provide a strategy for the further study of native monoamine transporters.

## Materials and Methods

### Antibody purification

The 15B8 Fab construct (50) was cloned into the pFastBac-dual vector, including a GP64 signal sequence. A mCherry tag, followed by twin Strep [TrpSerHisProGlnPheGluLys(GlyGlyGlySer)2GlyGlySerAlaTrpSerHisProGlnPheGluLys] and His_10_ purification tags, were fused to the C-terminus of the heavy chain. Baculovirus was prepared according to standard methods. The Sf9 cells were infected by the recombinant baculovirus at a cell density of 2 × 10^6^ ml^−1^ at 27°C. The culture supernatant was then collected 96 h after infection by centrifugation at 5,000 rpm for 20 min using a JLA 8.1000 rotor at 4°C. The 15B8 Fab was purified from Sf9 supernatant by metal ion affinity chromatography followed by size exclusion chromatography.

### Isolation of native pSERT

One porcine brain (∼150 g), obtained from a local slaughterhouse, was homogenized with a Dounce homogenizer in ice-cold Tris-buffered saline buffer (TBS: 20 mM Tris-HCl, pH 8.0, 150 mM NaCl) supplemented with 1 mM phenyl methylsulfonyl fluoride (PMSF), 0.8 µM aprotinin, 2 µg ml^−1^ leupeptin, and 2 µM pepstatin. The homogenized brain suspension was then sonicated using a sonicator equipped with a tip size of 1.27 cm, for 15 min with 3 s on and 5 s off, at medium power, on ice. The resulting solution was clarified by centrifugation for 20 min at 10,000 g at 4°C, the supernatant was collected and applied for further centrifugation at 40,000 rpm for 1 h at 4°C (45 Ti fixed-angle rotor, Beckman) to pellet the membranes. The membranes were resuspended in 40 ml ice-cold TBS and further homogenized with a Dounce homogenizer. The membranes were solubilized in 100 ml ice-cold TBS containing 20 mM n-dodecyl-β-D-maltoside (DDM) and 2.5 mM cholesteryl hemisuccinate (CHS) in the presence of 1 mg of 15B8 Fab, 100 µM methamphetamine or 10 µM cocaine, for 1 h at 4°C. The lysate was centrifuged at 40,000 rpm for 50 min at 4°C (45 Ti fixed-angle rotor, Beckman) and the transporter-Fab complex was isolated by affinity chromatography using Strep-Tactin resin. The complex was further purified by fluorescence-detection size-exclusion chromatography (FSEC) (71) on a Superose 6 Increase 10/300 column in a buffer composed of 20 mM Tris-HCl (pH 8) supplemented with 100 mM NaCl, 1 mM DDM, 0.2 mM CHS, and 100 µM methamphetamine or 10 µM cocaine. The peak fraction containing the native pSERT-Fab complexes was manually collected and used for biochemical and single particle cryo-EM analysis.

### Screening solubilization conditions

Porcine brain membranes were prepared as described above. The membrane fraction was resuspended and solubilized in ice-cold TBS containing buffer I (20 mM DDM, 2.5 mM CHS), buffer II (1% Digitonin), buffer III (1% Amphipol 17), or buffer IV (1% SMA- XIRAN30010), respectively. Mammalian cells expressing recombinant hSERT were solubilized with buffer I for comparison, as previously described (72). All the solubilizations were performed in the presence of 1 uM paroxetine for 1 h at 4°C. The solubilized solutions were clarified by centrifuged at 40,000 rpm using a TLA-55 rotor for 20 min at 4°C. The supernatant was then examined by FSEC on a Superose 6 Increase 10/300 column.

### Western blot analysis

Purified native pSERT was run on a SDS-PAGE gel and subsequently transferred to a nitrocellulose membrane. Antibodies used for detection were 10F2, a monoclonal antibody generated in house. An IRDye 680RD anti-mouse secondary antibody (LI-COR), was used for visualization. Blots were developed from the secondary antibody at a ratio of 1:10,000 and imaged by Odyssey® DLx Imaging System.

### Radioligand binding assay

A saturation binding experiment using [^3^H]paroxetine was performed via the scintillation proximity assay (SPA) (73) using the lysate of porcine brain membranes in SPA buffer (20 mM Tris-HCl, pH 8, 100 mM NaCl, 1 mM DDM, 0.2 mM CHS). The membrane lysates were mixed with Cu-YSi beads (0.5 mg ml^−1^) in SPA buffer, and [^3^H]paroxetine at a concentration of 0.3 to 40 nM. Nonspecific binding was estimated by experiments that included 100 μM cold S-citalopram. Data were analyzed using a single-site binding function.

Methamphetamine and cocaine competition binding experiments were performed using SPA with Cu-YSi beads (0.5 mg ml^−1^) in SPA buffer. For the methamphetamine competition assays, SPA was performed with Strep-purified native pSERT, 10 nM [^3^H]paroxetine, and 1 μM to 100 mM cold methamphetamine. For the cocaine competition assays, SPA was done with Strep-purified native pSERT and 10 nM [^3^H]paroxetine in the presence of 1 nM to 1 mM cold cocaine. Experiments were performed in triplicate. The error bars for each data point represent the standard error of the mean (SEM). K_i_ values were determined with the Cheng-Prusoff equation (74) in GraphPad Prism.

### Cryo-EM sample preparation and data acquisition

The purified native pSERT-15B8 Fab complex was concentrated to 0.1 mg ml^−1^, after which either 10 mM methamphetamine or 1 mM cocaine, together with 100 μM fluorinated n-octyl-β-D-maltoside (final concentration) were added prior to grid preparation. A droplet of 2.5 μl of protein solution was applied to glow-discharged Quantifoil 200 or 300 mesh 2/1 or 1.2/1.3 gold grids covered by 2 nm of continuous carbon film. The grids were blotted for 2.0 s at 100% humidity at 20°C, followed by plunging into liquid ethane cooled by liquid nitrogen, using a Vitrobot Mark IV. The native pSERT datasets were collected on a 300 kV FEI Titan Krios microscope located at the HHMI Janelia Research Campus, equipped with a K3 detector, at a nominal magnification of 105,000x, corresponding to a pixel size of 0.831 Å. The typical defocus values ranged from −1.0 to −2.5 μm. Each stack was exposed for 4.0 s and dose-fractionated into 60 frames, with a total dose of 60 e^−^ Å^−2^. Images were recorded using the automated acquisition program SerialEM (75).

### Cryo-EM image processing

The beam-induced motion was corrected by MotionCor2 (76). The defocus values were estimated by Gctf (77) and particles were picked by blob-picker in cryoSPARC (78). We explored blob picking with elliptical blobs of sufficient sizes to capture pSERT oligomers yet after two rounds of 2D classification, we found only pSERT monomers, and proceeded to select 2D classes with clear secondary structures. An initial model was generated by cryoSPARC and employed for heterogeneous refinement. Next, a round of 3D classification without image alignment was performed in RELION-3.1 (79), with a soft mask excluding the constant domain of 15B8 Fab and micelle. The selected particles were imported back to cryoSPARC for homogeneous refinement, local contrast transfer function (CTF) refinement, and nonuniform refinement. The local resolution of the final map was estimated in cryoSPARC.

For the pSERT-15B8 Fab complex in the presence of (+)-methamphetamine, 7,794,907 particles were picked in cryoSPARC, which after rounds of 2D classification and heterogeneous refinement, left 348,745 particles (binned to a 200-pixel box, 1.662 Å pixel^−1^). Particles were reextracted (360-pixel box, 0.831 Å pixel^−1^) and subjected for homogeneous refinement and nonuniform refinement in cryoSPARC, then subjected to 3D classification with 10 classes in RELION-3.1 without image alignment (360-pixel box, 0.831 Å pixel^−1^). Particles from three classes with clear TM features were combined and subjected to homogeneous refinement, local CTF refinement, and nonuniform refinement in cryoSPARC, respectively (*SI Appendix*, Fig. S2).

For the pSERT-15B8 Fab complex with cocaine, a total of 7,560,137 particles were picked from 16,094 movies in cryoSPARC with a box size of 200 pixels (1.662 Å pixel^−1^). After rounds of 2D classification and heterogeneous refinement, 338,343 particles were selected, reextracted (400-pixel box, 0.831 Å pixel^−1^), and subjected to homogeneous refinement, nonuniform refinement in cryoSPARC, and further subjected to 3D classification with 10 classes in RELION-3.1 without image alignment. Two well-resolved classes with 243,207 particles were combined and further refined in cryoSPARC with homogeneous refinement, local CTF refinement, and nonuniform refinement (*SI Appendix,* Fig. S3).

### Model building and refinement

Interpretation of the cryo-EM maps exploited rigid-body fitting of the SERT-antibody complex models derived from previous cryo-EM studies. The outward-open ΔN72/C13 SERT-15B8 Fab complex with a 5-HT model (PDB code: 7LIA) was used as a reference. The initial model was generated via rigid-body fitting of the homology models to the density map in UCSF ChimeraX (80). The model was then manually adjusted in COOT (81). The model was further refined using real-space refinement in PHENIX (82). Strikingly, for both complexes there was no density for the amino and carboxy terminal and thus the protein models begin at residue 116 and end at residue 654. Careful scrutiny of density maps showed evidence for *N-*linked glycosylation at Asn 245 but no evidence for phosphorylation or other post translational modification, at the present resolution. Figures were prepared in UCSF ChimeraX.

### System preparation for MD simulations

We performed molecular dynamics (MD) simulations of the cocaine-pSERT complex in a hydrated lipid bilayer to explore the identity of the ligand in the allosteric site ligand. Triplicates of five simulation systems were performed, with the allosteric site occupied by either docosahexaenoic acid (DHA) in charged or neutral forms (DHA^−^ or DHA^0^, respectively), cholesterol (CHOL), cholesteryl hemisuccinate (CHS), or n-dodecyl-β-D-maltoside (DDM). The initial coordinates of CHS, DHA^−^, and DDM were transferred from the experimental models, while CHOL was constructed into the CHS model, and DHA^0^ was constructed by protonating the DHA^−^ model using the PSFGEN plugin in VMD (83). The pSERT protein model contains residues 116-654. The missing side chains and hydrogen atoms in the protein were added using the PSFGEN plugin in VMD (83). The co-crystalized antibody fragment was removed. All bound Na^+^ and Cl^−^ ions, the cocaine molecule, and the two cholesterol molecules bound to transmembrane helices were retained. Glu173 was modeled as a protonated side chain according to pK_a_ calculations in PROPKA 3.0 (84). A disulfide bond was introduced between Cys 237 and Cys 246. Neutral N-terminal and C-terminal ‘caps’ were added to the first and last residues of the protein segment, respectively. All protein models were internally hydrated using the DOWSER plugin (85, 86) of VMD and externally solvated using the SOLVATE program (https://www.mpinat.mpg.de/grubmueller/solvate). The models were then oriented according to the Orientations of Proteins in Membranes (OPM) database (87) and embedded in a lipid bilayer consisting of 218 1-palmitoyl-2-oleoyl-sn-glycero-3-phosphocholine (POPC) and 94 CHOL molecules from CHARMM-GUI (88). The systems were next solvated and neutralized with a 150 mM NaCl aqueous solution in VMD (83), resulting in simulation systems of ∼160,000 atoms, with approximate dimensions of 112 Å × 112 Å × 120 Å before equilibration. Each simulation system was replicated into three independent copies with lipid distribution randomized by shuffling lipid molecules within each leaflet using the VMD plugin Membrane Mixer (89).

### Equilibrium MD simulations

All simulations were performed using NAMD2 (90, 91) and the CHARMM36m force fields (91) for proteins, CHARMM36 force fields for lipids (including CHS, CHOL, DHA^−^ and DHA^0^) and detergent DDM (92), and the TIP3P model for water (93), along with the NBFIX modifications for non-bonded interactions (94, 95). The force field parameters for cocaine were obtained from the CGenFF server (96). All simulations were carried out as isothermal-isobaric (NPT) ensembles under periodic boundary conditions and simulated in a flexible cell, whose dimensions could change independently while keeping a constant ratio in the *xy* (membrane) plane. A constant temperature of 310 K was maintained using Langevin dynamics with a 1.0-ps^-1^ damping coefficient, and a constant pressure of 1.01325 bar was maintained with the Langevin piston Nosé-Hoover method (97). van der Waals non-bonded interactions were calculated with a 12-Å cut-off, and a switching function applied between 10 Å and 12 Å. Long-range, electrostatic non-bonded interactions were calculated with the particle mesh Ewald (PME) method (98). Bond lengths involving hydrogen atoms were constrained using the SHAKE (99) and SETTLE algorithms (100). Simulations were integrated in 2 fs time steps, and trajectories were recorded every 10 ps.

The five simulation systems, each replicated into three independent copies, were simulated following these steps: (1) 3,000 steps of energy minimization; (2) 15 ns of MD equilibration, during which Cα atoms of the protein, non-hydrogen atoms of ligands, and all bound ions were restrained by harmonic potentials with progressively decreasing force constants (k = 5, 2.5, 1 kcal mol^-1^ Å^-2^ for 5 ns each) to allow for protein side chain relaxation and protein hydration; (3) 150 ns MD run, during which harmonic potentials (k = 1 kcal mol^-1^ Å^-2^) were applied to only Cα atoms to avoid undesired protein conformational deviation but allowing free diffusion of the allosteric site ligand; (4) 150 ns production MD run without restraints.

### Free energy characterization of ligand binding

Bias-exchange umbrella sampling (BEUS) method (101) was employed to characterize the binding energy profiles of CHOL, DHA^−^, and DDM to the allosteric site. The ligand- TM1b/TM6a distance, measured as the center-of-mass distance between the non-hydrogen atoms in the ligand and Cα atoms in the extracellular ends of TM1b and TM6a (residues 145- 148 and 361-364), was chosen as the reaction coordinate to sample ligand binding. The initial distances for the modeled CHOL, DHA^−^, and DDM were 17.2, 15.2, and 13.9 Å, respectively. We chose reaction coordinates ranging from 14 to 21.5 Å for CHOL, 15 to 22.5 Å and DHA^−^, and 13 to 20.5 Å for DDM, to sample their unbinding from the allosteric site. Each reaction coordinate was divided into 16 windows with a spacing of 0.5 Å. The initial conformations in each window (collectively shown in Fig. 4h as line representations) were captured from steered MD simulations using the COLVAR module^98^ in NAMD, in which the ligand was pulled away from the extracellular ends TM1b and TM6a (residues 145-148 and 361-364) to the desired distances using a harmonic potential (k = 10 kcal mol^-1^ Å^-2^) moving at a 0.5 Å ns^-1^ rate. The BEUS simulations were performed for 60 ns in each window. The Hamiltonian replica exchange was attempted every 1 ps between neighboring windows. Weighted Histogram Analysis Method (WHAM) (102, 103) was used to construct the free energy profiles and perform error analysis using the Monte Carlo bootstrapping method.

### Data sharing plan

The 3D cryo-EM density maps and molecular coordinates have been deposited in the Electron Microscopy Data Bank (EMDB) and Protein Data Bank (PDB) for the SERT-15B8- Fab-methamphetamine outward (EMD-27384; PDB 8DE4) and SERT-15B8-Fab-cocaine outward-occluded (EMD-27383; PDB 8DE3) reconstructions and structures, respectively.

## Acknowledgments

We thank Rui Yan at the HHMI Janelia CryoEM Facility for help in microscope operation and data collection, critiques from the Biophysics Colab by Azadeh Shahsavar and Steffen Sinning, as well as input from the reviewers of a previous version of the manuscript submitted to *Nature Comm*. A portion of this research was supported by NIH grant U24GM129547 and performed at the PNCC at OHSU and accessed through EMSL (grid.436923.9), a DOE Office of Science User Facility sponsored by the Office of Biological and Environmental Research. Simulations in this study have been performed using allocations at National Science Foundation Supercomputing Centers (XSEDE grant MCA06N060). This work was otherwise funded by the NIH (R01 MH070039, P41 GM104601 and R24 GM145965). E.G. is supported by Jennifer and Bernard LaCroute and is an investigator of the Howard Hughes Medical Institute.

## Contacts

Dongxue Yang; Zhiyu Zhao; Emad Tajkhorshid; Eric Gouaux

