## Supplementary material for "Structures and membrane interactions of native serotonin transporter in complexes with psychostimulants": SI Appendix

*Eric Gouaux

**This PDF file includes:**

Figures S1 to S8
Table S1

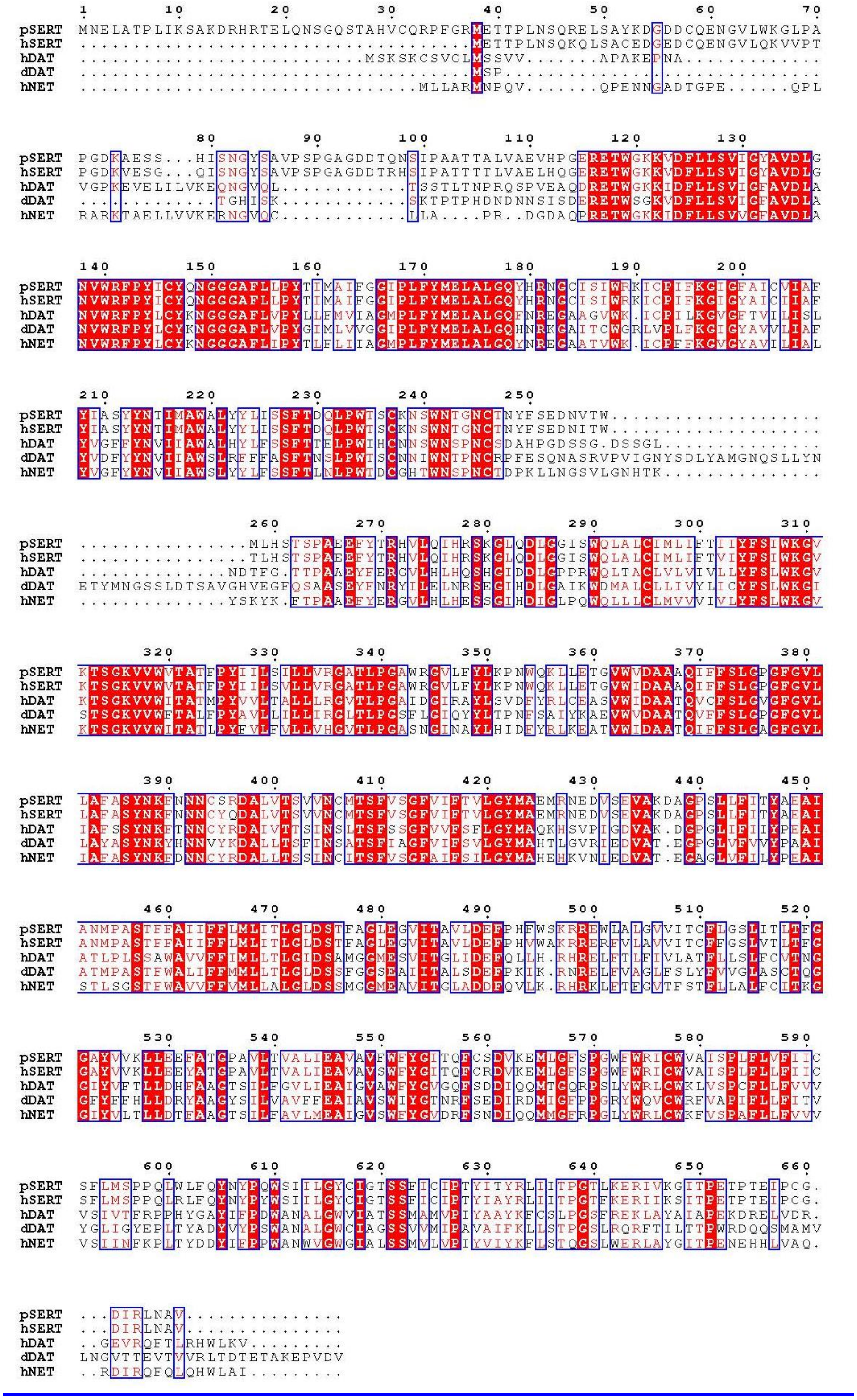

**Figure S1. Multiple sequence alignment of SERT, DAT and NET orthologues.** Residues in black are not conserved, those in red are conservatively substituted, and those in white with red background are conserved.

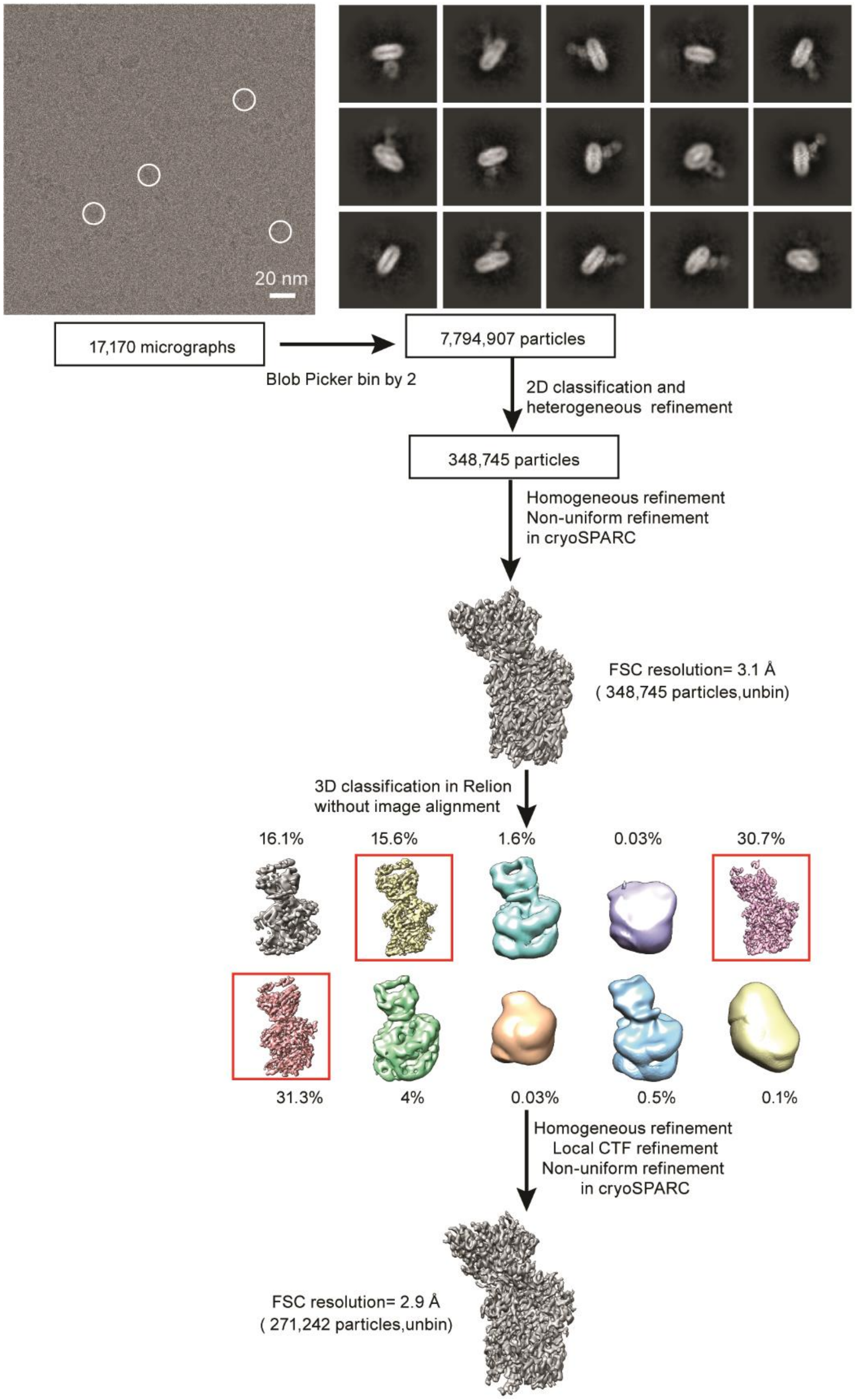

**Figure S2. 3D reconstruction of the pSERT-methamphetamine complex.** Flow chart for cryo-EM data analysis.

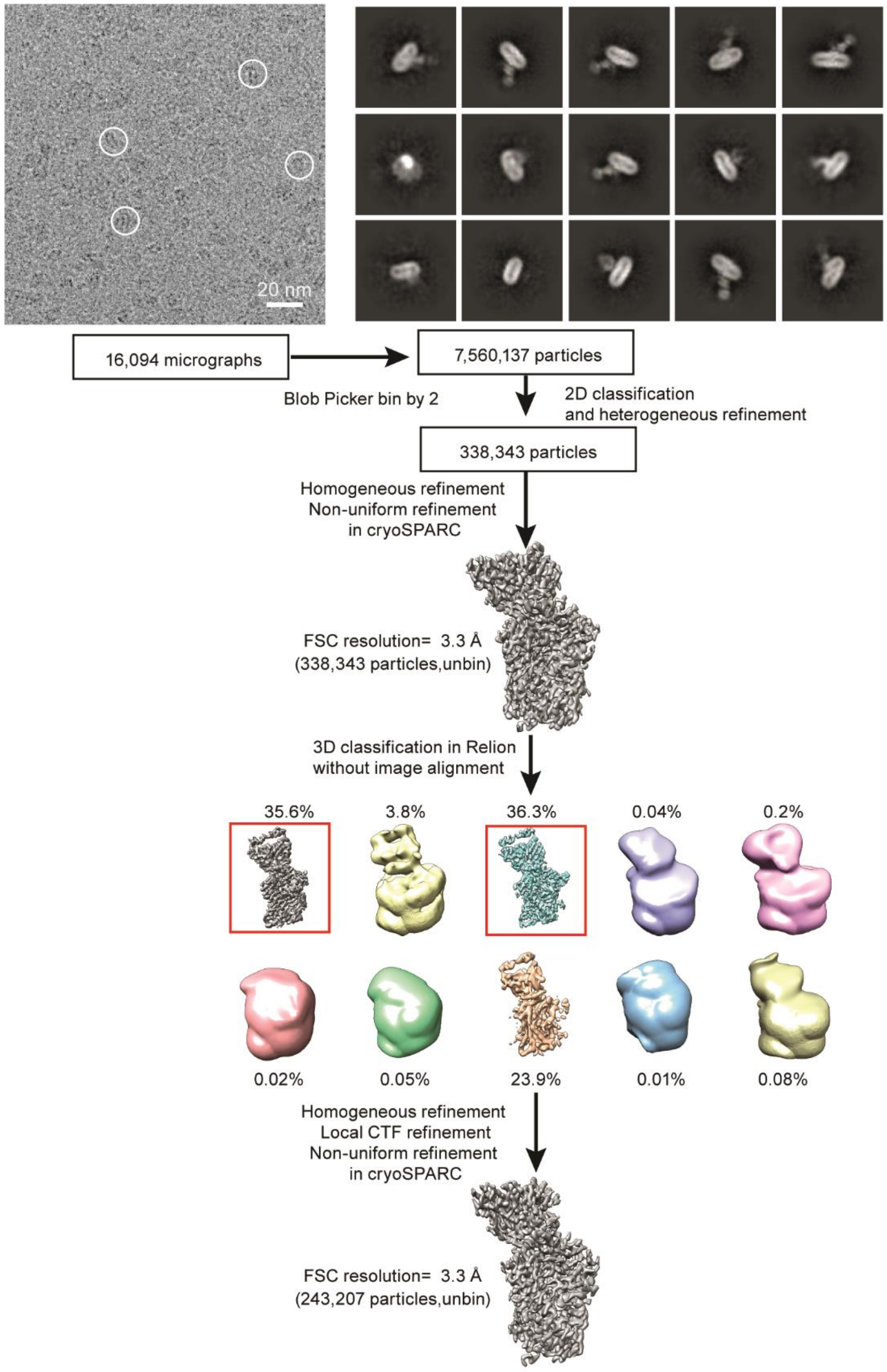

**Figure S3. Cryo-EM analysis of pSERT-cocaine complex.** Flow chart for cryo-EM data analysis.

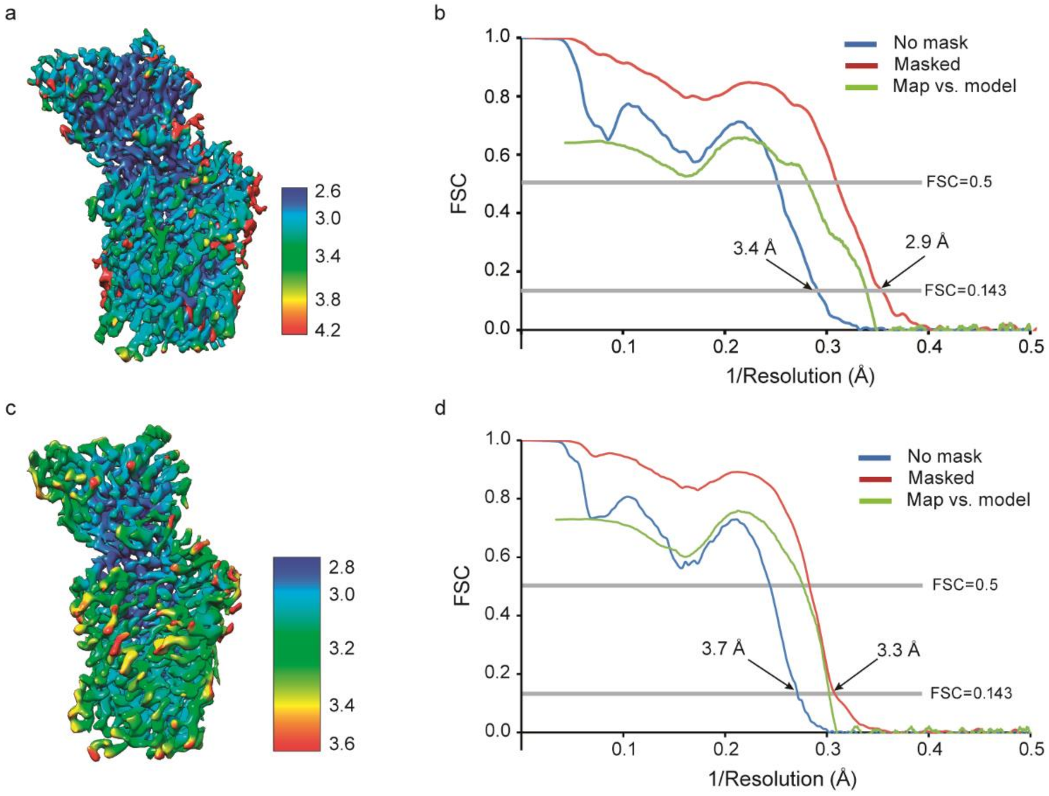

**Figure S4. Local resolution maps, overall plotted resolutions, and global map-model agreements. a** (+)-methamphetamine-pSERT Fab complex map is colored by local resolution. **b** Fourier shell correlation (FSC) curves for (+)-methamphetamine-pSERT Fab complex. **c** Local-resolution distribution of the cocaine-pSERT Fab map. **d** Map-map and map-model FSC curves for cocaine-pSERT Fab complex.

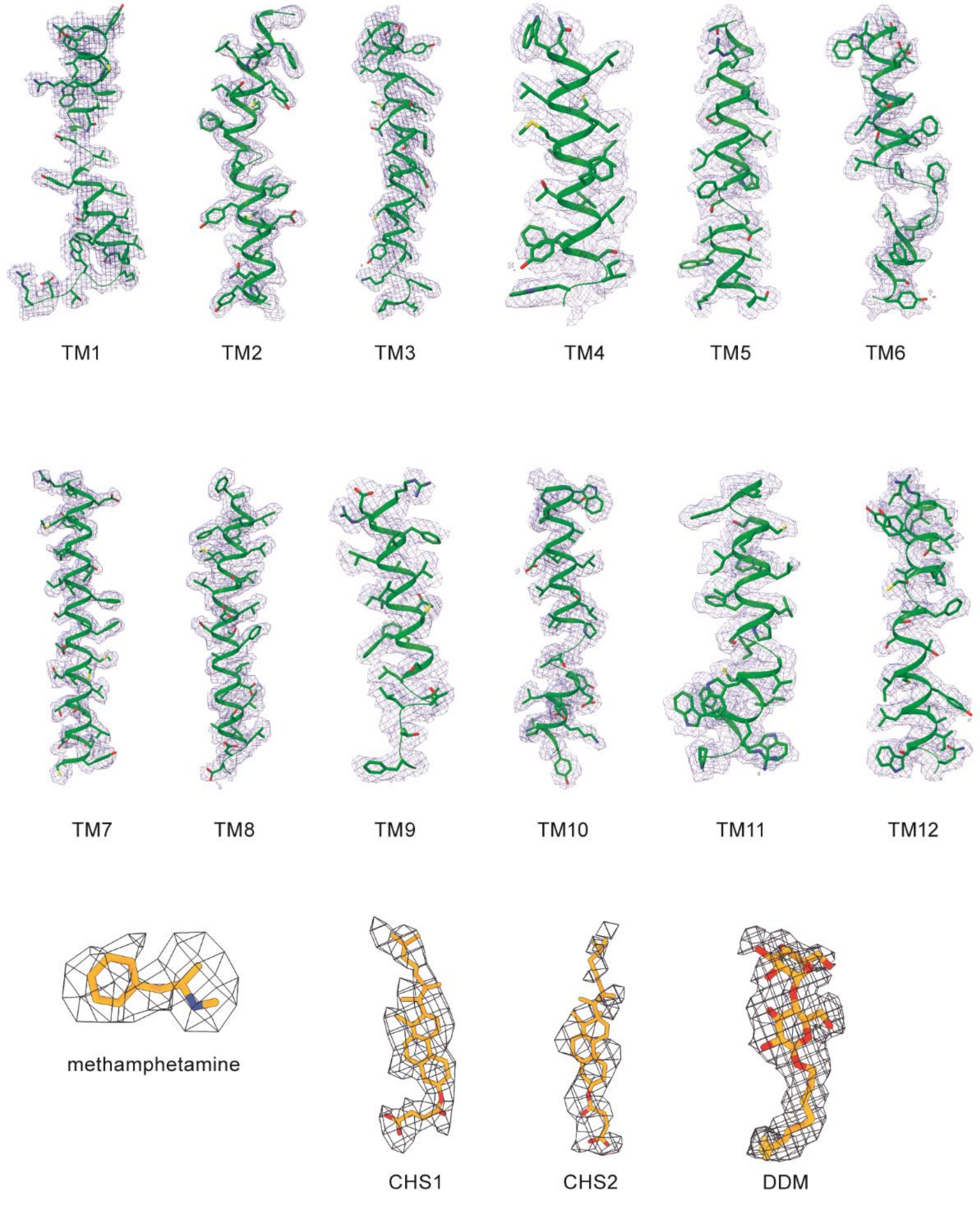

**Figure S5. Representative densities of (+)-methamphetamine-pSERT Fab complex.** Representative densities for transmembrane helices, (+)-methamphetamine cholesteryl hemisuccinate (CHS), and dodecylmaltoside (DDM).

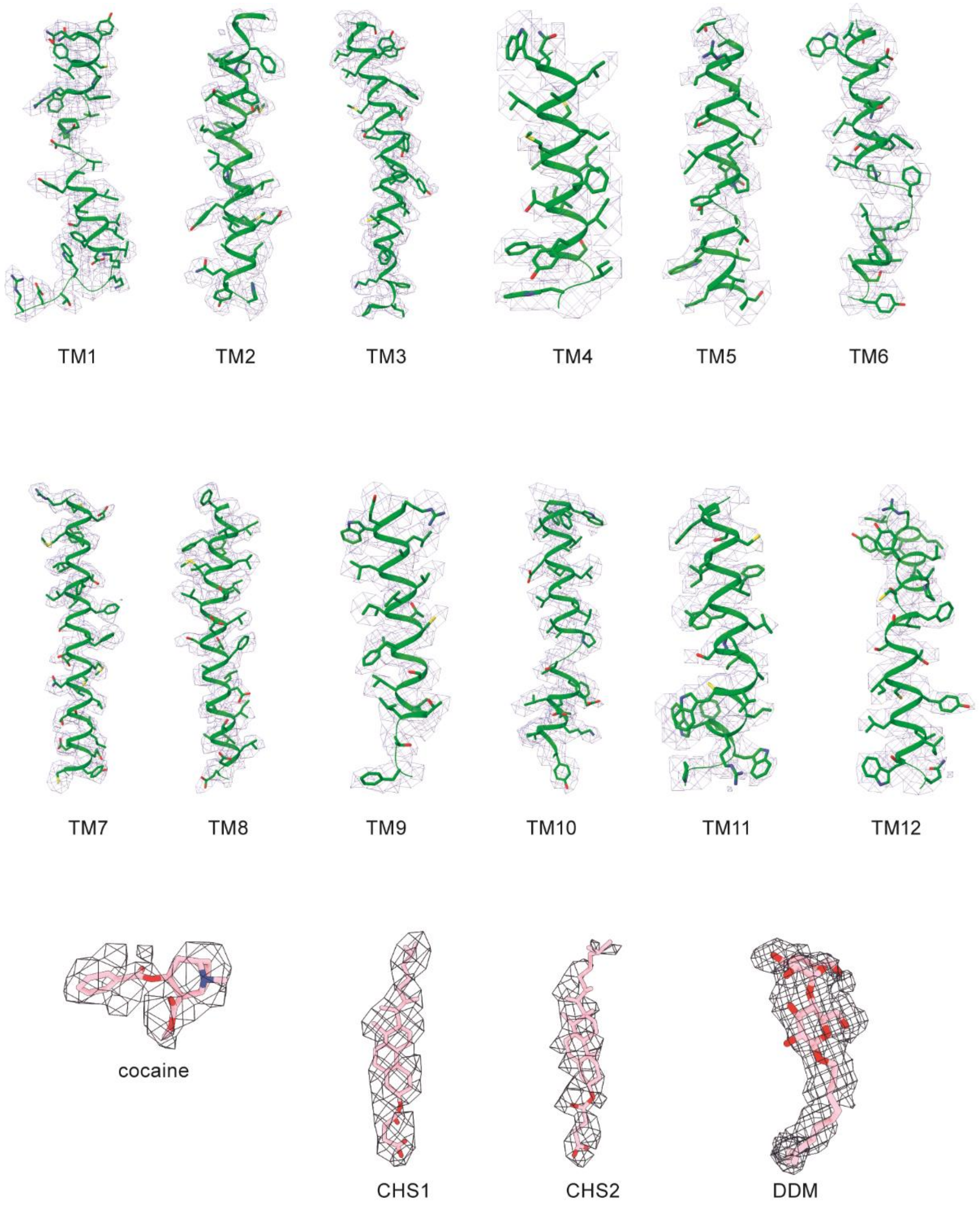

**Figure S6. Representative densities of cocaine-pSERT Fab complex.** Density fitting of transmembrane helices, cocaine, CHS, and DDM.

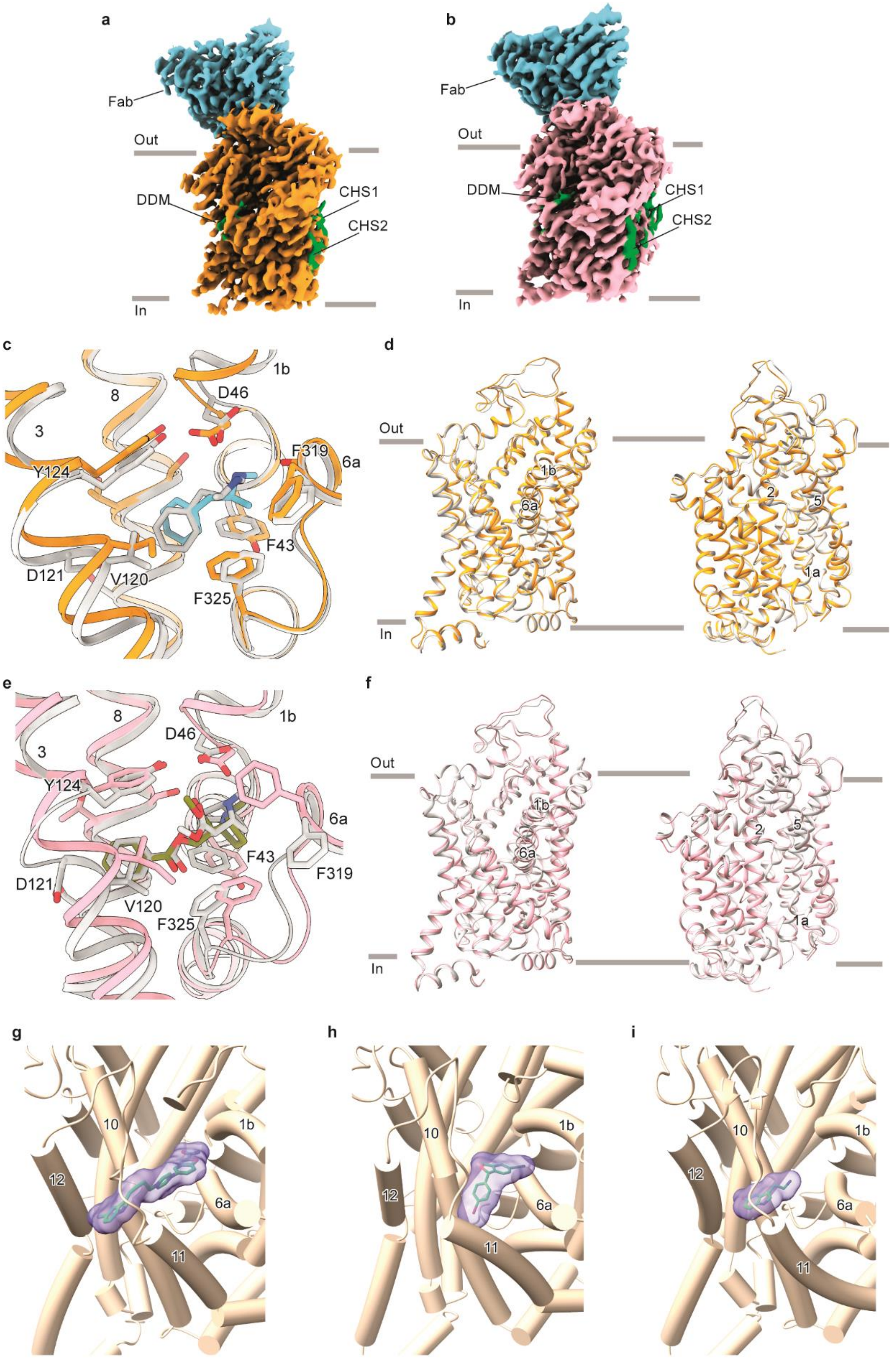

a
Fab
Out
In
DDM
CHS1
CHS2
b
Fab
Out
In
DDM
CHS1
CHS2
c
8
1b
3
D46
F319
6a
Y124
F43
D121
V120
F325
d
Out
In
1b
6a
2
5
1a
e
8
1b
3
D46
Y124
6a
F43
F319
D121
V120
F325
f
Out
In
1b
6a
2
5
1a
g
10
1b
12
6a
11
h
10
1b
12
6a
11
i
10
1b
12
6a
11

**Figure S7. The cryo-EM density maps of ligand-bound pSERT Fab complex and comparison of ligand binding in the central sites of SERT with dDAT. a** and **b** The cryo-EM density maps of (+)-methamphetamine-pSERT (a) and cocaine-pSERT (b) complex. **c** Superposition of the binding pockets of the (+)-methamphetamine-dDAT structure (PDB code: 4XP6) in grey with binding pockets of (+)-methamphetamine-pSERT (orange). Residues interacting with (+)-methamphetamine in dDAT have been indicated. **d** The superimposed pSERT-(+)-methamphetamine and hSERT-5-HT (outward-facing, PDB code: 7LIA). **e** Superposition of the central binding pocket of cocaine-dDAT structure (PDB code: 4XP4, grey), with central binding site of cocaine-pSERT (pink). Residues interacting with cocaine in dDAT have been indicated. **f** Structural comparison of pSERT-cocaine and hSERT-5-HT (outward-facing, PDB code: 7LIA). **g-h** Occupancy of the allosteric site by vilazodone (g), citalopram (h), and serotonin (i).

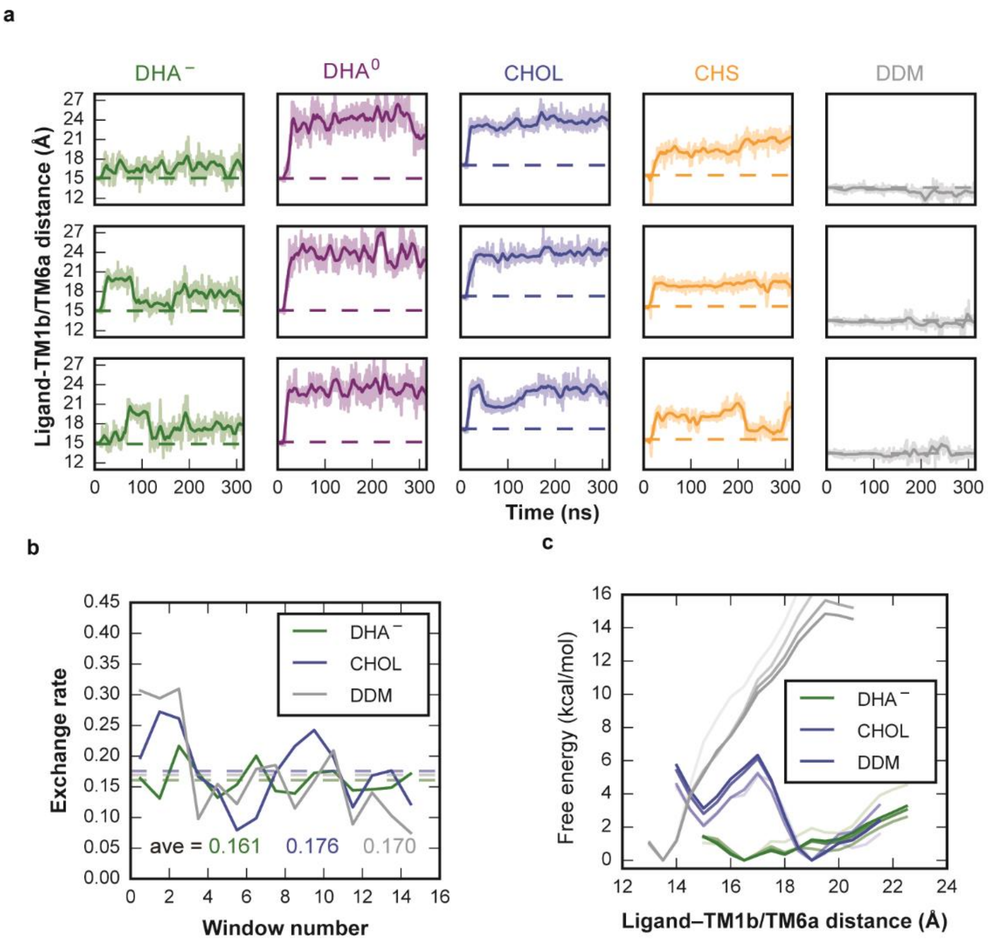

**Figure S8. Stability of lipids in the allosteric site monitored by root-mean-square deviation (RMSD) and convergence of BEUS simulations. a** Time series of ligand-TM1b/TM6a distance of the ligands at the allosteric site. Data for DHA⁻, DHA0, CHOL, CHS, and DDM are plotted in green, purple, blue, orange, and gray, respectively, and are shown for the three independent simulations in each case. The ligand-TM1b/TM6a distance is measured as the center-of-mass distance between heavy atoms in the ligand and Cα atoms from TM1b and TM6a (residues 145-148 and 361-364). Dashed lines indicate the initial distances of each molecule. Plots are smoothed using a sliding window of 1 ns. **b** Exchange rates between neighboring windows monitored during the 60-ns BEUS simulations. **c** Convergence of the free energy profiles along the increase in time (10, 30, 50, and 60 ns, indicated by increasing opacities) of the simulations.

**Table S1. Supplementary Table 1 Cryo-EM data collection, refinement and validation statistics**

| | **SERT-methamphetamine** (EMD-) (PDB ) | **SERT-cocaine** (EMD-) (PDB) |
|---|---|---|
| **Data collection and processing** | | |
| **Magnification** | 105,000 | |
| **Voltage (kV)** | 300 | |
| **Electron exposure (e–/Å$^2$)** | 60 | |
| **Defocus range (μm)** | -1.0 to -2.5 | |
| **Pixel size (Å)** | 0.831 | |
| **Symmetry imposed** | C1 | C1 |
| **Initial particle images (no.)** | 7,794,907 | 7,560,137 |
| **Final particle images (no.)** | 271,242 | 243,207 |
| **Map resolution (Å)**<br>**FSC threshold** | 2.9<br>0.143 | 3.3<br>0.143 |
| **Map resolution range (Å)** | 4.2-2.6 | 3.6-2.8 |
| **Refinement** | | |
| **Initial model used (PDB code)** | 7LIA | 7LIA |
| **Initial model CC**<br>**Model resolution (Å)**<br>**FSC threshold** | 0.628<br>3.4<br>0.5 | 0.623<br>3.6<br>0.5 |
| **Map sharpening *B* factor (Å$^2$)** | -95.3 | -103.1 |
| **Model composition** | | |
| **Non-hydrogen atoms**<br>**Protein residues**<br>**Ligands (atoms)** | 6188<br>769<br>116 | 6199<br>769<br>127 |
| ***B* factors (Å$^2$)** | | |
| **Protein**<br>**Ligand** | 59<br>77 | 29<br>36 |
| **R.m.s. deviations** | | |
| **Bond lengths (Å)**<br>**Bond angles (°)** | 0.002<br>0.631 | 0.002<br>0.505 |
| **Validation** | | |
| **Refined model CC**<br>**MolProbity score**<br>**Clashscore**<br>**Poor rotamers (%)** | 0.680<br>2.15<br>8.76<br>4.35 | 0.684<br>1.55<br>6.79<br>0.00 |
| **Ramachandran plot** | | |
| **Favored (%)**<br>**Allowed (%)**<br>**Disallowed (%)** | 96.85<br>3.15<br>0 | 96.97<br>3.03<br>0 |
